# Wild rice (*O. latifolia*) from natural ecosystems in the Pantanal region of Brazil: host to *Fusarium incarnatum-equiseti* species complex and highly contaminated by zearalenone

**DOI:** 10.1101/2020.07.06.190306

**Authors:** Sabina Moser Tralamazza, Karim Cristina Piacentini, Geovana Dagostim Savi, Lorena Carnielli-Queiroz, Lívia de Carvalho Fontes, Camila Siedlarczyk Martins, Benedito Corrêa, Liliana Oliveira Rocha

## Abstract

We assessed the mycobiota diversity and mycotoxin levels present in wild rice (*Oryza latifolia*) from the Pantanal region of Brazil; fundamental aspects of which are severely understudied as an edible plant from a natural ecosystem. We found a variety of fungal species contaminating the rice samples; the most frequent genera being *Fusarium, Nigrospora* and *Cladosporium* (35.9%, 26.1% and 15%, respectively). Within the *Fusarium* genus, the wild rice samples were mostly contaminated by the *Fusarium incarnatum-equiseti* species complex (FIESC) (80%) along with *Fusarium fujikuroi* species complex (20%). Phylogenetic analysis supported multiple FIESC species and gave strong support to the presence of two previously uncharacterized lineages within the complex (LN1 and LN2). Deoxynivalenol (DON) and zearalenone (ZEA) chemical analysis showed that most of the isolates were DON/ZEA producers and some were defined as high ZEA producers, displaying abundant ZEA levels over DON (over 19 times more). Suggesting that ZEA likely has a key adaptive role for FIESC in wild rice (*O. latifolia*). Mycotoxin determination in the rice samples revealed high frequency of ZEA, and 85% of rice samples had levels >100 μg/kg; the recommended limit set by regulatory agencies. DON was only detected in 5.2% of the samples. Our data shows that FIESC species are the main source of ZEA contamination in wild rice and the excessive levels of ZEA found in the rice samples raises considerable safety concerns regarding wild rice consumption by humans and animals.

## 1 Introduction

The Pantanal region is a 140,000 km^2^ sedimentary floodplain in western Brazil and one of the largest wetlands in the world (Pott and Silva, 2015); which experiences months-long floods every year during the rainy season from October to April (Bergier and Assine, 2016). The region harbors more than 200 wild grass species (Pott and Pott, 2000) that are commonly used for cattle grazing and are also a food source for native wildlife (Pott and Pott, 2004).

*Oryza latifolia* is a tetraploid wild species of rice, with a distribution ranging from Mexico to Brazil and the Caribbean Islands (Tateoka, 1962). The species is characterized as drought resistant, aquatic emergent and is largely found in the Pantanal wetland of Brazil (Bertazonni and Alves Damasceno-Júnior, 2011).

*O. latifolia* has been employed as a genetic resource to improve resistance to biotic and abiotic stress in conventional rice crops (*O. sativa*). Notable examples include resistance to bacterial blight, the brown planthopper (*Nilaparvata lugens*) and white-backed planthopper (*Sogatella furcifera*) (Multani et al., 2003, Angeles-Shim et al., 2020). More importantly, wild rice is also a source of nutrition for local communities (Bertazonni and Alves Damasceno-Júnior, 2011, Bortolloto et al., 2017), forage for livestock (Pott & Pott, 2000) and a component of wild animal diets, like jaguars, pumas and ocelots (Montalvo et al., 2020).

Despite being a food source for humans and animals, fundamental aspects of food-safety, such as the microbial diversity, and the presence of hazardous toxins, are severely understudied in wild rice from natural ecosystems. The lack of information is worrisome as a multitude of studies have shown that rice can be heavily afflicted by fungal pathogens in the field, particularly mycotoxigenic species of the *Fusarium* genus (Petrovic et al., 2013, Gonçalves et al., 2019). Their presence can cause significant economic losses through crop diseases and production of hazardous toxins (mycotoxins) that hinders cereal commercialization as food and feedstuff (Brown and Proctor, 2013).

The *Fusarium fujikuroi* species complex (FFSC) is one of the most prominent *Fusarium* complexes in rice crops. The group which includes the species *F. fujikuroi, F. proliferatum* and *F. verticillioides* are reported as the causal agent of the fast-emerging Bakanae disease. This disease can cause seedling blight, root and crown rot, etiolation, and the excessive elongation of infected rice plants. (Gupta et al., 2015). The FFSC members are also prolific producers of fumonisin, a mycotoxin which can have carcinogenic, hepatotoxic, nephrotoxic and embryotoxic effects in laboratory animals. In humans fumonisin is associated with esophageal cancer and neural tube defects (Scott, 2012).

Species within the *Fusarium graminearum* species complex (FGSC) became important plant pathogens in major rice producing regions, such as China (Qiu and Shi, 2014, Yang et al., 2018) and Brazil (Gomes et al., 2015, Moreira et al., 2020). The FGSC includes distinct species capable of causing Fusarium head blight in cereals and producing the sesquiterpene trichothecenes and the non-steroidal estrogenic mycotoxin zearalenone. (O’Donnel et al., 2004, Aoki et al., 2012).

Recently, the *Fusarium incarnatum-equiseti* species complex (FIESC) gained attention as a relevant mycotoxigenic contaminant of crops worldwide (Goswami et al., 2005, Castellá and Cabañes, 2014, Avila, et al., 2019). This complex has an intricate taxonomy (O’Donnel et al., 2012), and ongoing studies (Villani et al., 2016) are trying to resolve the species complex phylogeny. The complex was divided in two large clades, named *incarnatum* and *equiseti* (O’Donnell et al., 2009), which currently comprise more than 31 phylogenetically distinct species (O’Donnel et al., 2012, Villani et al., 2016). Like, the FGSC, the species of this group are known to produce significant amounts of trichothecenes and zearalenone and other mycotoxins such as equisetin, butenolide and fusarohromanone (Thrane, 1989, Kosiak et al., 2005, Goswami et al., 2008).

Deoxynivalenol (DON), the most prevalent variant of trichothecene is reported to inhibit protein synthesis by binding to the ribosome and causing anorexia, immune dysregulation as well as growth, reproductive, and teratogenic effects in mammals (Chen, Kistler and Ma, 2019). Zearalenone (ZEA) has been highly associated with significant changes in reproductive organs and fertility loss in animals (Kowalska et al., 2016). Also, the toxin has been found to induce the production of progesterone, estradiol, testosterone in the cell line H295R, indicating its potential as an endocrine-disruptive agent in humans (Frizzell, et al., 2011).

The presence of fungi and mycotoxins in wild rice is still poorly understood. Yet, the use of edible wild plants from natural ecosystems is a relevant ecological alternative resource to deforestation and monocultures (Bartollo et al., 2017). Moreover, the consumption of *O. latifolia* has been gaining more traction in recent years because of its higher nutritional value in comparison to *O. sativa* (Bertazzoni and Damasceno-Júnior, 2011). Due to the increasing relevance of wild cereal consumption, including *O. latifolia*, it is essential to investigate the safety concerns regarding the introduction of these novel sources of nutrition for human and animal use. This study aims to characterize the unexplored diversity of *Fusarium* in the wild rice O*. latifolia* from the Midwest Pantanal region of Brazil through investigation of the rice fungal community, their mycotoxigenic potential, the rice mycotoxin content and possible link between natural and managed rice systems.

## 2 Materials and methods

### 2.1 Sample collection and fungal isolation

The Brazilian Pantanal region is characterized by annual and pluri-annual flooding, forming distinct sub-regions; including the Pantanal of Paraguay River, with local flora and fauna adapted to the seasonal water level variations (Alho and Sabino, 2012). Random sampling was adopted in this study due to the irregular distribution of the plants throughout the river. A total of 50 wild plants (five samples per point at 10 randomly selected location points) were collected from the Paraguay river close to the city of Corumba (−19°00’33.01” S −57°39’11.99” W), Mato Grosso do Sul, Brazil (Figure 1), in June 2016. The rice grains were placed in PCNB-PPA medium (Leslie and Summerell, 2008) and incubated at 25° C for 7 days for fungal isolation. After the incubation period the fungal colonies were identified based on morphology using MEA (Malt Extract Agar) and CYA (Czapek Yeast Extract Agar) media (Pitt and Hocking, 2009) and molecular markers.

**Figure 1.**
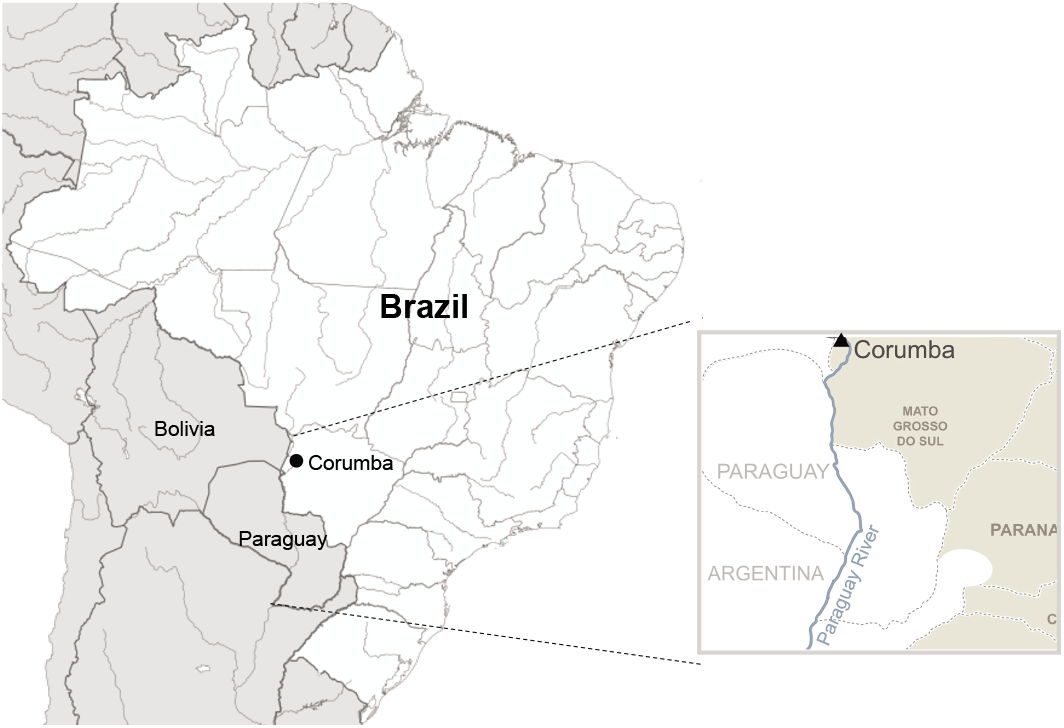
Map of Brazil indicating the site from which *Oryza latifolia* samples were sampled, Paraguay River, Corumba City (State of Mato Grosso do Sul, Brazil). Triangle marks the sampling area.

### 2.2 DNA extraction and PCR amplification

Fungal isolates were cultured on PDA medium for 5 days at 25° C. DNA extraction was conducted using the Easy-DNA kit (Invitrogen, Carlsbad, USA) according to manufacturer instructions. Genus level identification was carried out with the amplification of the partial sequence of the internal transcribed spacer (ITS) using primers set ITS1 and ITS2 (White, et al.,1990). Further identification of *Fusarium* isolates was conducted using the elongation factor (EF-1α) loci with primer set EF-1 (5’ ATGGGTAAGGARGACAAGAC 3’) and EF-2 (5’ GGARGTACCAGTSATCATGTT 3’) according to O’Donnell et al. (1998) protocol. DNA sequences were determined using ABI 3730 DNA Analyzer (Applied Biosystems, Foster City, USA) in the Human Genome and Stem Cell Research Center (HUG-CELL) (Sao Paulo, Brazil). The EF-1α sequences were deposited in the NCBI database (Supplementary Table S1).

### 2.3 Phylogenetic analysis

The resulting EF-1α sequences were aligned with ClustalW v2.1 (Thompson, 1994) plugin using Geneious v.11 software. The isolates within the FIESC were chosen in addition to several reference strains. *Fusarium chlamydosporum* strains (MRC117 and MRC35) were used as outgroup, based on the phylogenetic analysis performed by O’Donnel et al. (2018). The phylogenetic analysis was run on PAUP 4.0b10 (Swofford, 2002). The most parsimonious tree was inferred based on a heuristic search option with 1000 random additional sequences and tree-bisection-reconnection algorithm for branch swapping. JModelTest (Posada, 2008) was used to determine the best substitution model. We used Neighbour-Joining analysis and assessed clade stability using Maximum Parsimony Bootstrap Proportions (MPBS) with 1000 heuristic search replications with random sequence addition. We used Bayesian Likelihood analysis to generate Bayesian Posterior Probabilities (BPP) for consensus nodes using Mr Bayes 3.1 run with a 2,000,000-generation Monte Carlo Markov chain method with a burn-in of 500,000 trees. The phylogenetic trees were visualized using FigTree v.1.4 (University of Edinburgh, Edinburgh, United Kingdom).

### 2.4 Mycotoxin analysis

#### 2.4.1 Rice samples

The content analysis of DON and ZEA was assessed in 38 samples of wild rice according to Savi et al. (2018). Briefly, 2 g of ground rice was homogenized in 8 mL of acetonitrile:water:formic acid (80:19.9:0.1 v/v/v) and shaken for 60 min at 130 rpm. The mixture was centrifuged for 10 min at 3500 rpm. The resulting supernatant was dried in an amber vessel using a heat block and air stream at 60°C.

#### 2.4.2 Mycotoxigenic potential of FIESC strains

A total of 18 strains from the FIESC were selected and tested for their ability to produce DON and ZEA. To assess mycotoxin production the strains were cultured on PDA medium (three agar plugs, 6 mm in diameter) for 20 days at 24° C and 90% humidity for DON analysis (Savi et al., 2013b) and at 15° C and 80% humidity for ZEA analysis (Savi et al., 2013a). The grown cultures were transferred into Schott bottles with 30 mL of chloroform and shaken for 60 min for mycotoxin extraction, followed by filtration through anhydrous sodium sulfate (Na2SO4), the procedure was conducted three times. The extract was filtered with a hydrophilic PVDF membrane (0.22 μm) followed by evaporation using a heat block and air stream at 60°C. The residue was dissolved in 500 μL of mobile phase, consisting of 70% of water:methanol:acetic acid (94:5:1, v/v/v) and 30% of water:methanol:acetic acid (2:97:1, v/v/v). The extract (5 μL) was injected into the LC/MS-MS system (Savi et al., 2018).

#### 2.4.4 Chromatography conditions

The detection and quantification of DON and ZEA were carried out according to Savi et al. (2018) protocol. The analysis were performed in a LC/MS-MS system from Thermo Scientific^®^ (Bremen, Germany) composed of an ACCELA 600 quaternary pump, an ACCELAAS auto-sampler and a triple quadrupole mass spectrometer TSQ Quantum Max Analytes were separated on a C8 Luna column Phenomenex (150×2.0 mm, length, and diameter, respectively) with particle size of 3 μm (Torrance, USA). Eluent A (water: methanol: acetic acid, 94:5:1, v/v/v) and eluent B (water: methanol: acetic acid, 2:97:1, v/v/v) were used as mobile phase. The gradient program was applied at a flow rate of 0.2 mL/min under the following conditions: 0–1 min 55% eluent B; 1–3 min 55–100% B; 3.01–7 min 100% B and 7.01–12 min 55% B. The total analytical run time was 7.5 min and the retention time was 2.19 min and 6.55 min for DON and ZEA, respectively.

The mass spectrometer ionization conditions were 208° C for capillary temperature, 338° C for vaporizer temperature, 4500 V for spray voltage and 60 bar for sheath gas pressure. For selectivity, the mass spectrometer was operated at MRM mode monitoring three transitions per analyte using a collision gas pressure of 1.7 mTorr and collision energy (CE) ranging from 11 to 40 eV. The mass spectrometric conditions were optimized (quantification transition: 203 m/z and confirmation transition: 175, 91 m/z for DON; quantification transition: 283 m/z and confirmation transition: 187, 185 m/z for ZEA) with reasonably high signal intensities in positive ESI mode (ESI+), and protonated molecules [M+H] (297 m/z for DON and 319 m/z for ZEA). All measurements were done with the following settings: cone voltage 17, 18 e 39 V e Tube Lens 71 V for DON and cone voltage 11, 25 e 20 V e Tube Lens 79 V for ZEA.

#### 2.4.5 Validation of the method

To validate the method for extraction of mycotoxins in the rice grains and the fungal mycelia we follow the Commission Regulation guidelines (EC, 2000). Samples with non-detectable levels of mycotoxins were submitted to spiking experiments to determine the limit of detection (LOD), limit of quantification (LOQ), recovery, repeatability and selectivity/specificity. A six-point calibration curve was made with a mixture of DON and ZEA standards in the following concentrations: 0.025, 0.0375, 0.0625, 0.125, 0.375, 0.500 g/mL. To determine the LOD and LOQ, blank samples were fortified with different mycotoxin concentration levels and the experiments replicated on distinct days. The LOD was defined as the minimum concentration of an analyte in the spiked sample with a signal noise ratio equal to 3 and LOQ with a signal noise ratio equal to 10.

## 3 Results

### 3.1 Mycobiota diversity in wild rice

We investigated the fungal community present in wild rice (*O. latifolia*) from the Pantanal region of Brazil to determine diversity, mycotoxigenic potential and possible link between natural and managed rice systems. We found a variety of fungal species co-contaminating the rice samples; the most frequent genera being *Fusarium, Nigrospora* and *Cladosporium* (35.9%, 26.1% and 15%, respectively) (Figure 2). We performed a comparative sequence analysis of the *Fusarium* strains using NCBI blastn search engine, and based on the top alignment identity, we found the wild rice samples were mostly contaminated with species from the *Fusarium incarnatum-equiseti* species complex (80%) and the *Fusarium fujikuroi* species complex (20%) (Figure 2). Unexpectedly, we did not isolate any species from the FGSC. Next, due to the high frequency of FIESC isolates, we performed phylogenetic analysis using publicly available sequences of FIESC species as references to further resolve the FIESC population inhabiting wild rice of natural ecosystems.

**Figure 2.**
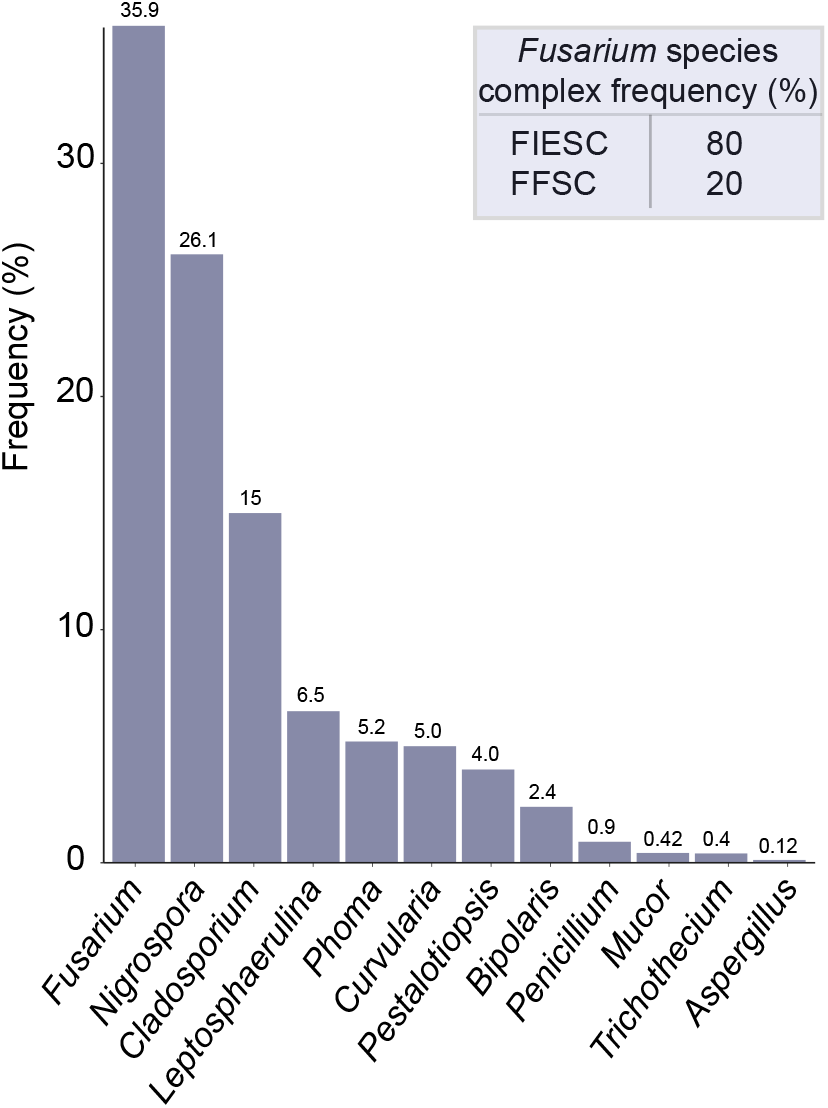
Frequency of fungal genera isolated from *O. latifolia* from natural ecosystems of the Brazilian Pantanal region. FIESC – *Fusarium incarnatum-equiseti* species complex, FFSC – *Fusarium fujikuroi* species complex.

### 3.2 Phylogenetic analysis of the FIESC strains

Tree topology based on the *EF-1* locus and supported with bootstrap and posterior probabilities showed that *O. latifolia* harbors a large group of phylogenetically distinct species within the FIESC (Figure 3). Our phylogenetic tree resolved all isolates within the *Fusarium incarnatum* clade. Part of the isolates grouped as FIESC15 (MS2763 and MS2965), FIESC16 (MS3369) and FIESC20 (MS2965) species. A single isolate (MS743) shared a monophyletic clade with FIESC25 and FIESC26. Interestingly, we also found two large groups that indicate two new lineages within the FIESC, here provisionally called LN1 and LN2. One of the new putative lineage (LN1) grouped closer to the species FIESC23 and the other lineage (LN2) shared a clade with the sister species FIESC24. (Figure 3).

**Figure 3.**
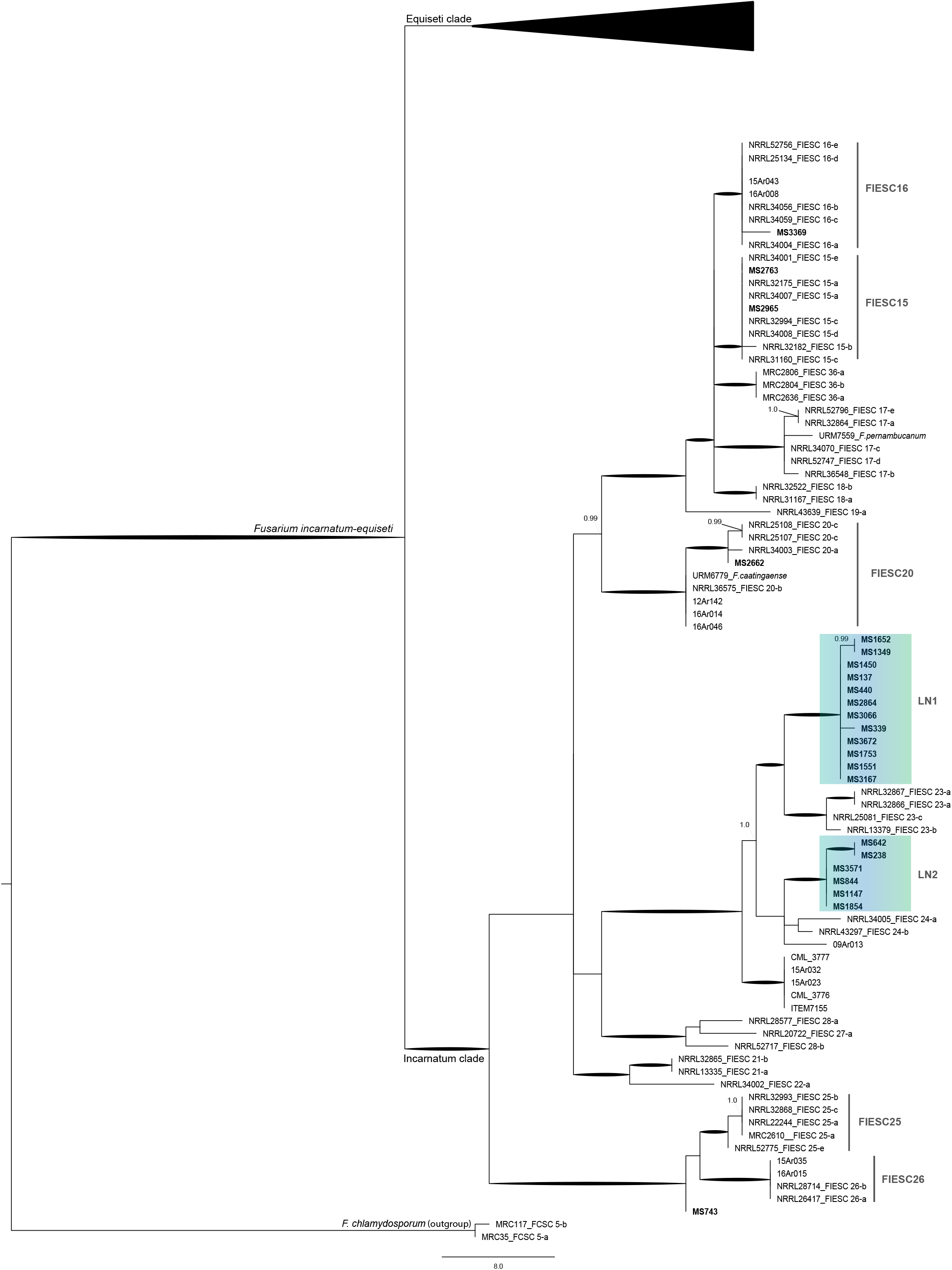
Maximum parsimony tree inferred from the EF-1α locus of the *Fusarium incarnatum-equiseti* species complex species (FIESC). Two strains of *F. chlamydosporum* included as outgroup based on O’Donnel et al. (2018). Bootstrap intervals (10,000 replications) >70% and Bayesian posterior probabilities >0.90 are indicated as branches in bold. Blue box highlights the putative new species within FIESC.

### 3.3 Toxigenic analysis of the FIESC strains

We assessed the toxigenic potential *in vitro* of the phylogenetically distinct FIESC strains to produce DON and ZEA. Most of the strains (88.8%) produced at least one type of mycotoxin. DON levels ranged from 13.5 to 41.0 μg/kg (mean of 23.4 μg/kg) and ZEA levels ranged from 7.5 to 757.6 μg/kg (mean of 123.2 μg/kg) (Figure 4A).

**Figure 4.**
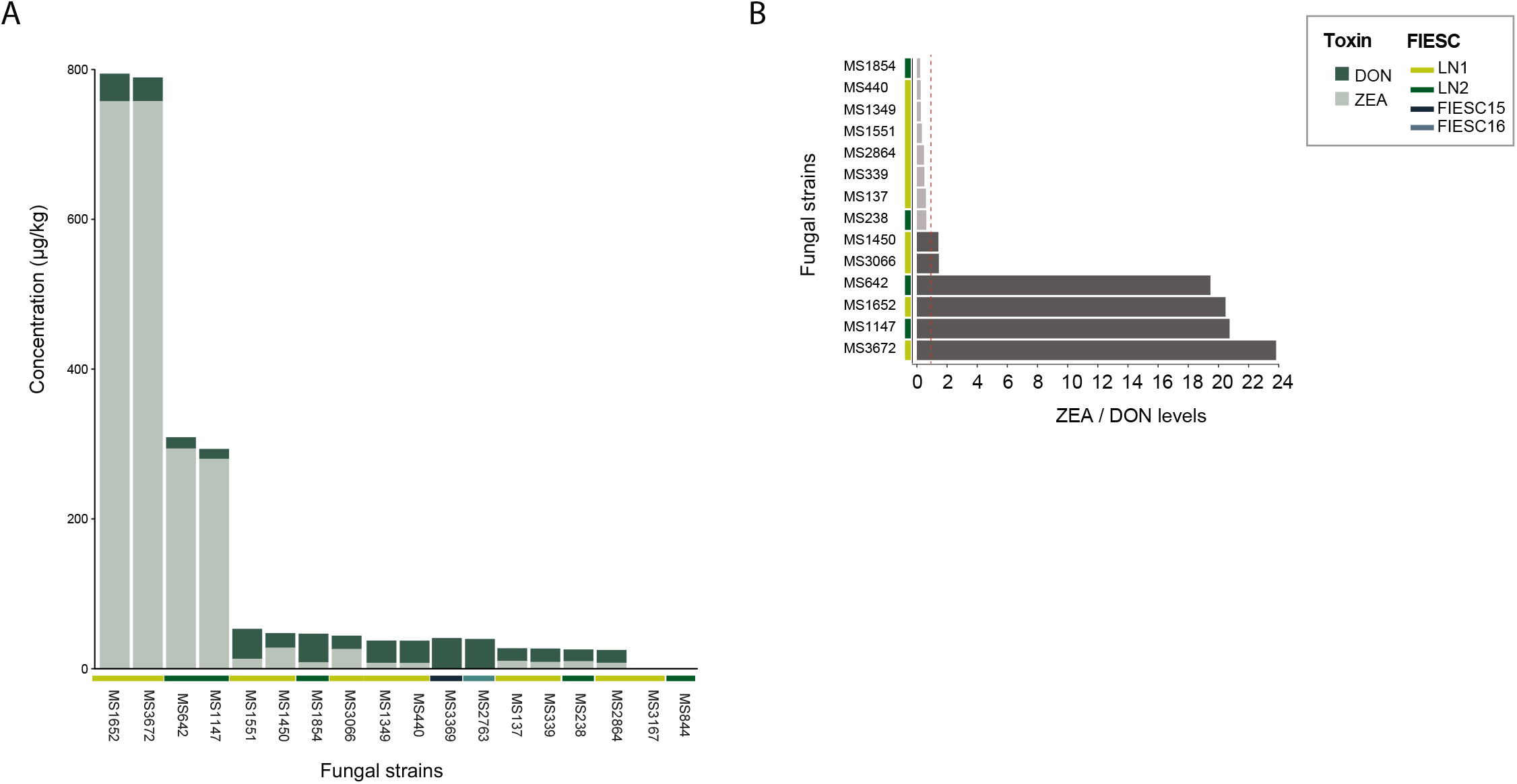
A – Concentration levels of deoxynivalenol (DON) and zearalenone (ZEA) produced by members of the *Fusarium incarnatum-equiseti* species complex *in vitro*. Figure 4. B – Ratio of ZEA over DON levels produced by the fungal strains. Dotted red line marks the ratio of one representing no difference.

The FIESC population of wild rice presented a diverse toxigenic profile. A great portion of the isolates (77.7%) produced both toxins and at relative similar rates (Figure 4). Two strains identified as FIESC15 (MS2769) and FIESC16 (MS3363) produced only DON at detectable levels. Interestingly, these strains are closer together and shared a large clade that includes FIESC15, 16, 17,18 and 36 (Figure 4).

We also found that 22.2% of the strains were high ZEA producers, displaying over 19 times more ZEA than DON levels (Figure 4B). All the major ZEA producers were part of the putative new lineages (LN1 and LN2) (Figure 4B, Supplementary Table 2). In two strains (MS3167 – LN1 and MS844 – LN2) no detectable levels of DON or ZEA were found. Although, the strains showed a diverse toxigenic profile, no clear relation was found between the toxin profile and the species phylogeny.

### 3.4 DON and ZEA analysis of the wild rice

We confirmed the presence of DON and ZEA in the wild rice samples. Only two samples were found to be contaminated by DON. These samples were co-contaminated with ZEA and displayed similar DON and ZEA contents (concentrations ranging from 81.7 to 92.5 ug/kg). Conversely, our analysis showed that most samples were highly contaminated by ZEA (92.1%), with levels ranging from 70.2 to 528.7 ug/kg (mean of 342.0 ug/kg) (Figure 5). Alarmingly, 85% of samples showed ZEA levels above 100 ug/kg (Figure 5, Supplementary Table 3), which is the maximum tolerated level specified by the European Commission for unprocessed cereals (other than maize) (EC, 2006).

**Figure 5.**
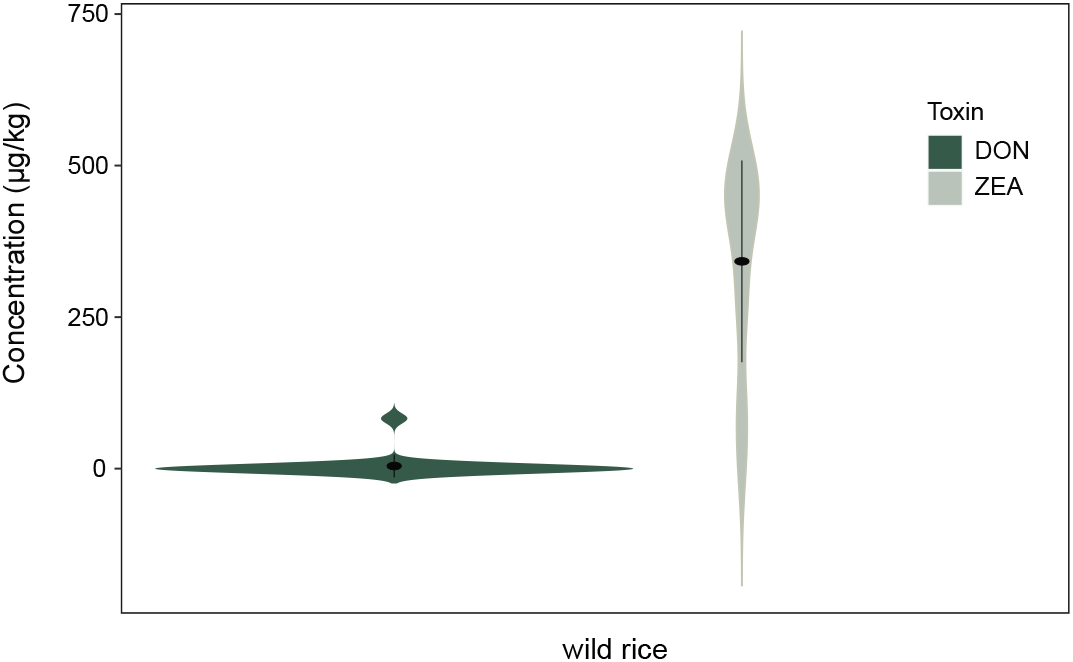
Deoxynivalenol (DON) and zearalenone (ZEA) content of wild rice (*O. latifolia*) from natural ecosystems of the Brazilian Pantanal region.

## 4 Discussion

We assessed the mycobiota diversity and mycotoxin levels present in the edible wild rice (*O. latifolia*) from the Pantanal region of Brazil. We also increased the currently available information of mycotoxin and fungal community contaminants on wild rice of natural ecosystems. Our work highlighted that *O. latifolia* harbors new lineages of the FIESC which are major ZEA producers. Our results also emphasized the importance of monitoring mycotoxins levels in alternative food sources.

### 4.1 Wild rice *O. latifolia* shares a similar mycobiota community with cultivated rice

Overall, the rice samples exhibited a similar mycobiota profile as previously reported in cultivated rice (Morillo et al., 2011, Ok et al., 2014, Katsurayama et al., 2020), where *Fusarium* spp. were the primary plant pathogen along with other fungi genera such as *Nigrospora*, *Cladosporium* and *Phoma*. Regarding wild rice, data is still scarce but a study on *Oryza australiensis*, a native wild rice of the Northern Territory of Australia, found a high presence of *Bypolaris oryzae* (the causal agent of brown spot), while *Fusarium* spp.*, Phoma* and *Cochliobolus* spp. were reported at lower frequencies (Pak et al., 2017). This difference could be explained by environment and host difference between studies.

Phylogenetic analysis revealed that the FIESC was a major contaminant of *O. latifolia*. While, no information is available concerning this specific rice species, other researchers analyzing the *Fusarium* community of *O. australiensis* identified the same species complexes and at analogous frequencies (FIESC - 55%, FFSC - 27 %, *F. longipes* – 14%) (Petrovic et al., 2013). Recently, Moreira et al. (2020) surveyed multiple regions of cultivated rice fields (*O. sativa*) in Brazil and reported FIESC as the most frequent *Fusarium* group of rice crops, followed by FFSC, FGSC and the *F. chlamydosporum* species complex across the country. Interestingly, they examined rice crops from Mato Grosso State, which is near the region where our samples were collected, and reported high infection with FIESC, followed by FFSC, and no presence of FGSC, which was congruent with our findings.

### 4.2 Wild rice harbors uncharacterized species of the FIESC

The challenging FIESC taxonomy (O’Donnel et al., 2012, Villani et al., 2016) makes the addition of strains from natural ecosystem hosts particularly relevant. Our phylogenetic analysis resolved all the isolated strains within the *Fusarium incarnatum* clade. We found a portion of the isolates grouped together with characterized FIESC species (FIESC15, FIESC16, FIESC20 and FIESC26) previously reported in cultivated rice (O’Donnel et al., 2012, Villani et al., 2016, Avila et al., 2020). Two isolates (MS2763 and MS2965) shared a monophyletic clade with FIESC15, a group with a wide range of hosts, having been associated to human infections (O’Donnel et al., 2009), insects (O’Donnel et al., 2012) and plants (Ramdial et a. 2016). To our knowledge this is the first time FIESC15 was described as a contaminant of rice grains. Interestingly, most of the analyzed strains formed two new lineages (LN1 and LN2) within the complex, which was supported with bootstrapping and posterior probability (LN1: 96.5% and 1.0; LN2: 94.9% and 1.0 for bootstrapping and posterior probability, respectively). The FIESC phylogenetic diversity is critically understudied and new species are continuously being described (Santos et al., 2019, Avila et al., 2019). Although, the evidence indicated new species, we believe the inclusion of more molecular markers (Summerell, 2019) will increase the confidence of these findings. According to recent genomic analysis performed with 13 FIESC strains, the group shares similar genome size (36.6 – 40 Mb) and gene content (12 −13k) but varies on the secondary metabolite repertoire (Villani et al., 2019) suggesting a possible adaptative function within the complex. However, information about aggressiveness, host range and geographical distribution of FIESC species is still lacking.

### 4.3 FIESC has a lead role in ZEA levels in the wild rice (*O. latifolia*)

The fungal toxigenic analysis shed light on important aspects of the species complex. FIESC15 and FIESC16 exclusively produced deoxynivalenol, which corroborates with a previous study where investigating the genomic diversity of 13 FIESC species reported that the zearalenone gene cluster is degenerated in FIESC15 (Villani et al., 2019). Currently, there is no available information about the gene cluster in FIESC16, nonetheless the FIESC15 and FIESC16 close relationship, could indicate the loss of a functional ZEA cluster in a recent common ancestor.

Most of the LN1 and LN2 strains produced DON and ZEA and some isolates were defined as high ZEA producers, displaying more than 19 times ZEA than DON levels. The strains belonging to the two new putative lineages were the most frequent isolates in the wild rice samples which could be a strong indication that ZEA has a key adaptative role for the group to inhabit wild rice (*O. latifolia*).

Zearalenone is a common contaminant of cereals (Tanaka et al., 2007) and it is usually found at relatively high frequencies in rice grains worldwide (40-60%) (Almeida et al., 2012, Savi et al, 2018, Golge and Kabak, 2020). Our data showed alarmingly high levels of ZEA in wild rice (>90%), with most of the samples exhibiting concentrations above the recommended limit (100 μg/kg) for unprocessed cereals (EU, 2006). ZEA contamination in rice crops and derived products have been associated to FGSC presence in the host (Savi et al., 2018, Ok et al., 2014). However, no species of the FGSC was isolated from *O. latifolia*. These findings along with the toxigenic ZEA profile of the strains strongly support that FIESC species are the main source of zearalenone contamination in *O. latifolia*. Our data corroborates previous hypotheses that in Brazil, high FGSC infections in rice systems are concentrated in small grain (e.g. wheat) producing regions, which may act as major hosts for FGSC species (Del Ponte et al., 2015, Moreira et al., 2020). Additionally, the concerning frequency and concentration levels of ZEA in the rice grains indicate that FIESC could be a much more relevant ZEA producer in Brazilian crops than previously contemplated.

We described previously uncharacterized FIESC members likely responsible for the elevated levels of zearalenone in *O. latifolia*, signifying a complex fungal diversity in wild rice from natural ecosystems. These findings give rise to many concerns since excessive levels of mycotoxins could greatly impair the safety of wild rice consumption for humans and animals. In addition, *O. latifolia* could act as a pathogen and/or a genetic pool reservoir and impact managed rice systems (Suproniene et al., 2019, Dong et al., 2020). *B. oryzae* strains isolated from the wild rice *O. australiensis* were reported as highly virulent to cultivated rice (*O. sativa*) of North Queensland, Australia (Pak et al., 2017). *Mycosphaerella graminicola*, a recent pathogen of domesticated wheat is an example of how the introduction of a new host rapidly selected a highly specialized pathogen from wild grasses close relatives (Stukenbrock et al., 2011). Nonetheless, our study highlights the importance to investigate fungal pathogens of wild hosts and how they could impact natural and managed systems.

## Acknowledgements

This research was supported by the São Paulo Research Foundation (FAPESP) grant processes 2015/21378-7 and 2016/04364-5.

## Notes

### Competing Interest Statement

The authors have declared no competing interest.

